# Modelling contractile ring formation and division to daughter cells for simulating proliferative multicellular dynamics

**DOI:** 10.1101/2023.03.26.534262

**Authors:** Satoru Okuda, Tetsuya Hiraiwa

## Abstract

Cell proliferation is a fundamental process underlying embryogenesis, homeostasis, wound healing, and cancer. The process involves multiple events during each cell cycle, such as cell growth, contractile ring formation, and division to daughter cells, which affect the surrounding cell population geometrically and mechanically. However, existing methods do not comprehensively describe the dynamics of multicellular structures involving cell proliferation at a subcellular resolution. In this study, we present a novel model for proliferative multicellular dynamics at the subcellular level by building upon the nonconservative fluid membrane (NCF) model that we developed in earlier research. The NCF model utilizes a dynamically-rearranging closed triangular mesh to depict the shape of each cell, enabling us to analyze cell dynamics over extended periods beyond each cell cycle, during which cell surface components undergo dynamic turnover. The proposed model represents the process of cell proliferation by incorporating cell volume growth and contractile ring formation through an energy function and topologically dividing each cell at the cleavage furrow formed by the ring. Numerical simulations demonstrated that the model recapitulated the process of cell proliferation at subcellular resolution, including cell volume growth, cleavage furrow formation, and division to daughter cells. Further analyses suggested that the orientation of actomyosin stress in the contractile ring plays a crucial role in the cleavage furrow formation, i.e., circumferential orientation can form a cleavage furrow but isotropic orientation cannot. Furthermore, the model replicated tissue-scale multicellular dynamics, where the successive proliferation of adhesive cells led to the formation of a cell sheet and stratification on the substrate. Overall, the proposed model provides a basis for analyzing proliferative multicellular dynamics at subcellular resolution.

## 1. Introduction

Cells undergo successive rounds of proliferation, wherein each cell undergoes an increase in its volume followed by division into daughter cells through the formation of a contractile ring. There are various regulations governing the behavior of cell proliferation; for example, the duration of the cell cycle varies widely across cell types [1]. Asymmetric cell division leads to generating geometrical and biochemical differences in daughter cells such as cell size and differentiation [2]. The direction of cell division is regulated according to various rules, such as being globally aligned according to surrounding biochemical gradients [3] or locally determined according to the mechanical environment [4]. These behaviors play key roles in a wide range of multicellular dynamics, including embryogenesis, homeostasis, wound healing, and carcinogenesis.

During the process of growth and division, proliferating cells generate various types of mechanical forces at the subcellular resolution, including a pushing force induced by cell rounding and a contractile force produced by the contractile ring [1,5]. These forces act on surrounding cells and are fed back to the proliferating cells themselves, regulating their growth rate, the direction of their division, and the resulting cell shape and configuration [4,6]. The mechanical interactions among cells are integrated to form the entire tissue structure. Therefore, to understand the integration of forces from cells to tissue, it is necessary to elucidate the mechanical influence and role of proliferating cell behaviors in multicellular dynamics.

Computational simulations are widely used to analyze the mechanical processes of multicellular dynamics, including cell proliferation. Several methods, such as vertex models [7–9] and cell membrane models [10–12] have been proposed for this purpose. However, most of these methods ignore the process of cleavage furrow formation by a contractile ring and describe the mitotic process by simply dividing each cellular object, as in vertex models [13,14], cell membrane models [15,16], and phase-field model [17]. Other methods, which focus on the mitotic mechanisms of a single cell, describe cytoskeletal dynamics leading to the formation of the cleavage furrow [18]. Thus, while conventional methods can be applied to either the entire process of multicellular dynamics or the detailed process of single-cell proliferation, there is a lack of methods that can comprehensively describe both processes, i.e., multicellular dynamics at the subcellular resolution.

When modeling multicellular dynamics involving cell proliferation at the subcellular resolution, a caveat that must be considered is the fact that cell proliferation occurs over a timescale of minutes to hours, during which the cell surface membranes and cortical cytoskeletons are dynamically turned over through endocytosis and exocytosis, polymerization, and depolymerization. To analyze such long-term cell dynamics at the subcellular resolution, a nonconservative fluid membrane (NCF) model has been developed [19]. In this model, each cell shape is expressed by a closed triangular mesh, and its dynamics are expressed by an overdamped equation of motion without mass conservation. Furthermore, the NCF model has been applied to whole-cell adhesion dynamics by constructing a continuum model of the interactions at the interface between membranes [20]. Thus, investigating the mechanical effects of proliferating cell behaviors on long-term multicellular dynamics is a challenging problem that requires expressing cell proliferation within the NCF model framework.

In this study, we aimed to analyze proliferative multicellular dynamics by modeling the mechanical process of cell proliferation based on the NCF model framework. In the proposed model, cell growth is expressed by increasing cell volume through an energy function, and the contractile ring is expressed by introducing active stress that leads to forming a cleavage furrow. In addition, cell division is expressed by topologically dividing a triangular mesh of each cell at the cleavage furrow. We performed numerical simulations of a single-cell proliferation to verify that the proposed model recapitulates cell volume growth, cleavage furrow formation, and division to daughter cells at the subcellular resolution. Furthermore, we performed numerical simulations of successive proliferation of adhesive cells on a substrate to clarify that the proposed model can be applied to complex multicellular dynamics at the tissue level. Based on these analyses, we discussed the applicability and future perspectives of the proposed model.

## 2. Brief review: Nonconservative fluid membrane (NCF) model

In this article, we construct the computational model for simulating long-term multicellular dynamics with successive cell proliferation. For this purpose, we implemented the cell proliferation model into the NCF model, which we proposed previously to describe long-term cell surface dynamics with turnover [19]. Before introducing the cell proliferation model, in this section, we review the NCF model briefly.

The NCF model describes fluidic and nonconservative features of the cell surface membrane, i.e., the dynamic flow and turnover of the membrane, including phospholipids and cortical actin. In this model, the cell surface membrane is discretized by a triangular mesh, which is a unifying element for describing the three-dimensional (3D) shape of a cell (Fig. 1a-b). By focusing on long-term cell behaviors over tens of minutes, inertial forces can be ignored. The governing equation of the membrane, whose velocity vector is represented by ***u***, is expressed as

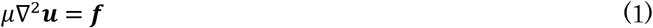

**Fig. 1:**
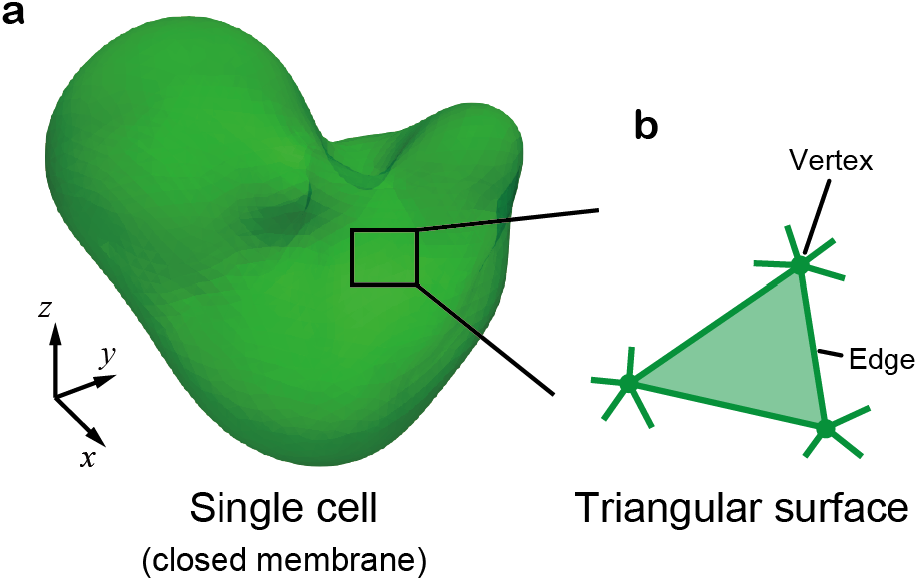
Discrete description of cell surface membrane in three dimensions. **a.** Single cell shape with a closed membrane. **b.** Triangular mesh describing cell shapes. Vertices and edges are shared with neighboring triangles. This figure has been modified from the previous study [19,20].

In Eq. (1), the left-hand term indicates viscous force per area on the membrane. The constant *μ* indicates the planar viscous coefficient of the cell surface membrane involving the cell cortex, i.e., the viscous coefficient in the planar direction of the membrane. In Eq. (1), the normal viscosity, i.e., the viscosity in the direction along membrane thickness is ignored, and the shear and dilational components of the planar viscous coefficients are assumed to have the same value. The right term, ***f***, represents the external force per area.

To discretize the forces in Eq. (1) on the cell surface into those on the vertices in the triangular mesh, the viscous force, *μ*▽^2^***u***, is discretized by the discrete type of the Laplace-Beltrami operator, and the external force, ***f***, is discretized by the gradients of an energy function, *U*, and dissipation function, *W*, as described in the previous study [19]. Therefore, Eq. (1) can be rewritten as

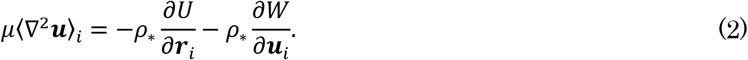

The left-hand side in Eq. (2) indicates the viscous force around the *i*th vertex. The first and second terms on the right-hand side indicate the energetic and viscous forces per area on the *i* th vertex, respectively. The constant *ρ*, is the mean numeric density of vertices on the membrane.

In Eq. (2), energetic forces involving active force that cells generate described by the function of *U*, such as the osmotic pressure, surface tension, bending moment, and cell-cell adhesion, are introduced. By representing the *i*th vertex location by *r_i_* (***u**_i_* = *∂**r**_i_*/*∂t*) and the active parameter by *a_i_* around the *i*th vertex, *U* is given as

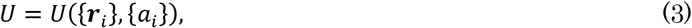

where { } indicate a set of values. Viscous forces on cell membrane from the environment, such as a viscous resistance from solvent and other cells, are described by the function *W*. Using ***r**_i_* and ***u**_i_*, *W* is given as

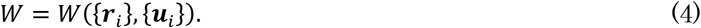

Equations (3) and (4) express approximate physical properties of cells around a non-equilibrium steady state.

During large deformations of cells, vertex distributions can be scattered by vertex displacements. To optimize the distribution, the mesh structure was dynamically rearranged using a dynamic remeshing algorithm that we previously proposed [19]. The algorithm comprises three steps: 1) the elimination and insertion of edges, 2) the regularization of edge connectivity, and 3) the centering of vertex locations. Geometric and energetic errors generated by this algorithm are negligible. The details of the NCF model are described in the previous work [19].

## 3. Modelling cell proliferation on the NCF model framework

In this section, we proposed a coarse-grained model that describes cell proliferation based on the NCF model framework (Fig. 2). The process of cell proliferation involves cell volume growth, contractile ring formation, and division to daughter cells. The proposed model describes this process by combining mechanical and topological operations, i.e., cell growth is expressed by an increase in cell volume through the energy function (Fig. 2a-b, Sec. 3.1). Actomyosin contraction of the contractile ring is expressed by introducing active stress through the energy function, leading to forming a cleavage furrow (Fig. 2b-c, Sec. 3.2). In addition, cell division is expressed by topologically dividing a triangular mesh of each cell at the cleavage furrow (Fig. 2c-d, Sec. 3.3).

**Fig. 2:**
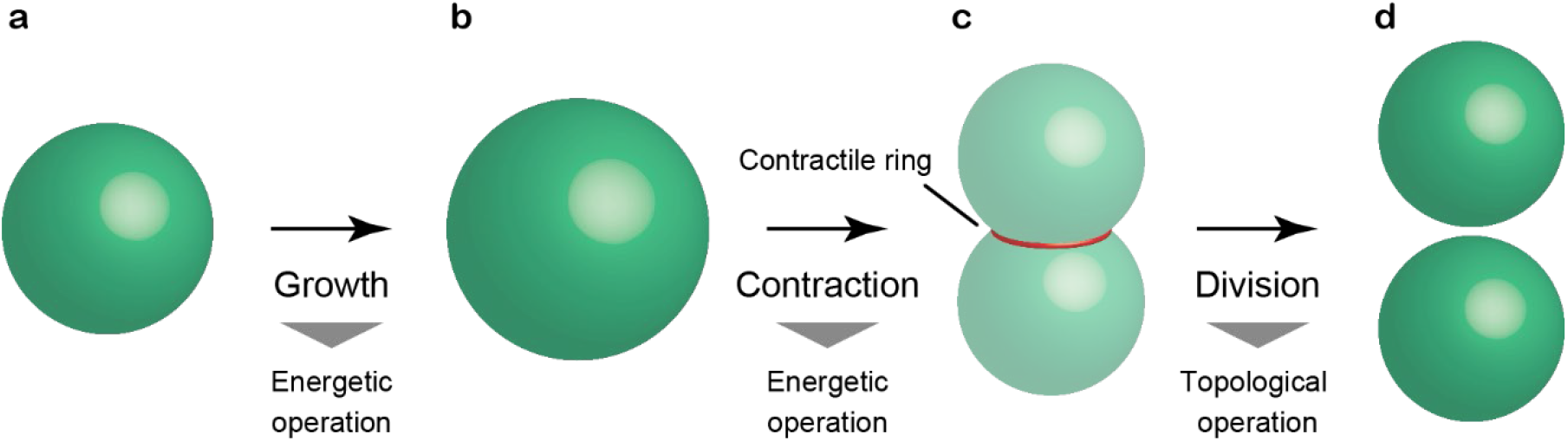
Description of cell proliferation process. Cell growth expressed as an increase in cell volume by the effective energy function (**a, b**). Cell contraction by a contractile ring expressed in terms of the effective energy function (**b, c**). Cell division expressed by topologically dividing the membrane into two parts (**c, d**).

### 3.1 Description of cell growth

Cell growth is expressed by an increase in cell volume through the energy function, *U*, in Eq. (3) (Fig. 2a-b). The energy function of the isolated *i*th cell, represented by 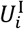, is given by

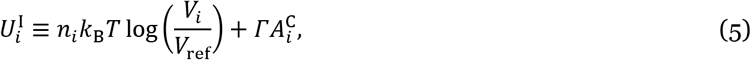

which has been used in the previous work of a 3D vertex model [21]. The first term on the right-hand side indicates osmotic energy, where constants *k_B_* and *T* are the Boltzmann constant and temperature. Variables *n_i_* and *V_i_* are the current number of intracellular molecules and the volume of the *i*th cell, respectively. Constant *V*_ref_ is the reference volume of cells. The second term on the right-hand side indicates surface energy, where constant *Γ* indicates a cell surface tension and variable 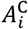 indicates the total surface area of the *i*th cell.

To express cell growth, for simplification, we assumed that the *i*th cell increases *n_i_* linearly with time from (2/3)*n*_ref_ to (4/3)*n*_ref_ and maintain it, where constant *n*_ref_ is the reference number of intracellular molecules.

After the growth, a contractile ring pinches off and divides the *i*th cell into two cells with the volume of (2/3)*n*_ref_.

Therefore, the time difference of *n_i_* can be given by

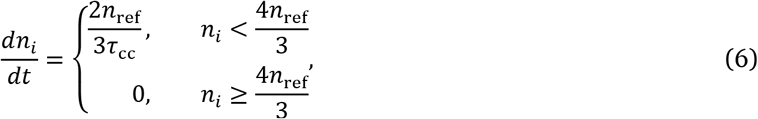

where constant *τ_cc_* is the cell cycle period.

Supposing that the isolated single cell is in equilibrium, the forces of the osmotic energy and surface energy balance within the *i*th cell, giving the relationship among physical parameters [21]. By regarding the number of intracellular molecules of *n*_ref_ as that of the cell with the volume of *V*_ref_, *n*_ref_ is given by

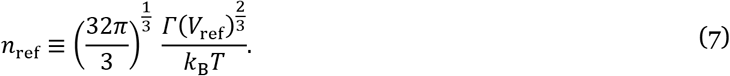

By considering *n_i_k*_B_*T* as a variable in Eq. (5), *n_i_k*_B_*T* is derived by Eqs. (6) and (7) with the constants *Γ* and *V*_ref_ as governing parameters.

### 3.2 Description of contractile ring formation

After cell growth, a contractile ring is formed around the center of the cell, leading to forming of a cleavage furrow (Fig. 2b-c). The location and orientation of the contractile ring of the *i*th cell are determined by introducing a division plane, which is defined as a plane that passes through the position represented by 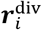, and has the normal vector represented by 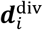 (Fig. 3a). For simplification, in this study, 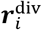 is given as a location vector of the cell centroid, and 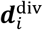 is given as a direction vector between a pair of vertices which has the longest length within the cell. Notably, regulations of cell division asymmetry and direction can be expressed by modifying the definitions of 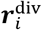 and 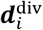, respectively.

**Fig. 3:**
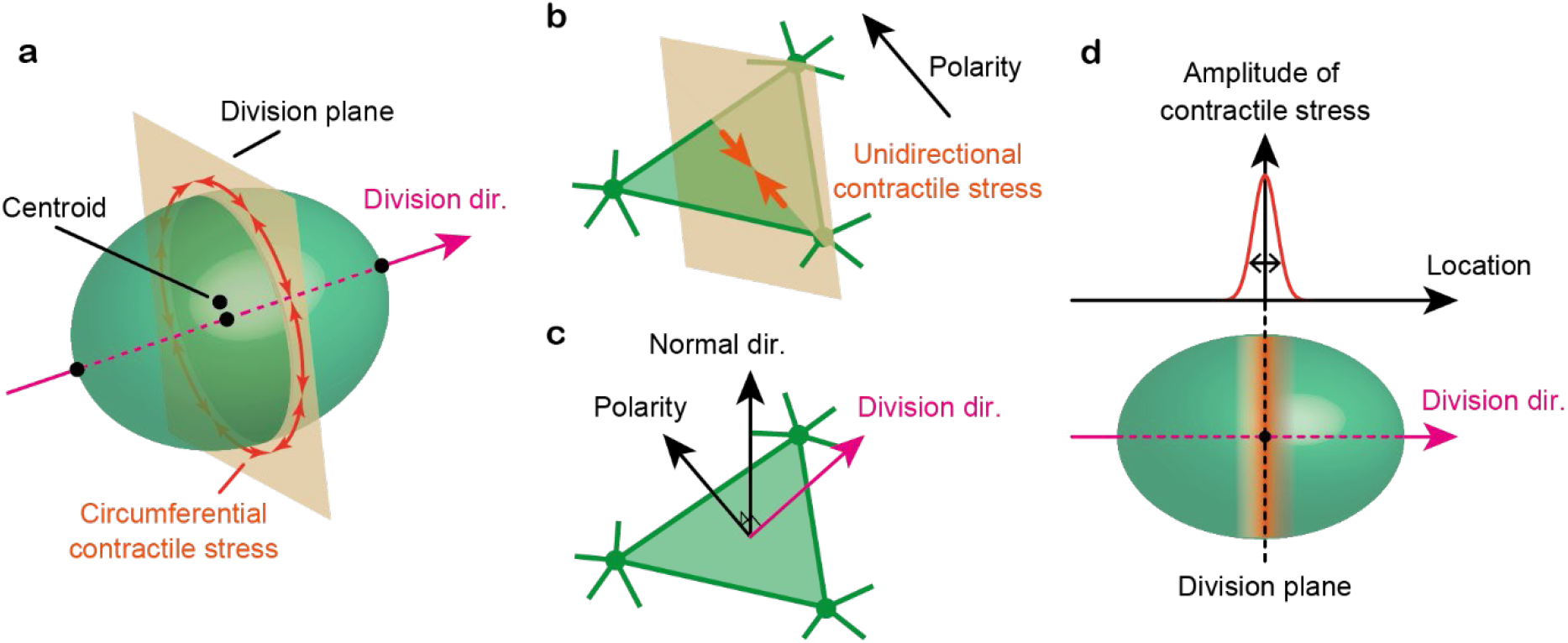
Description of contractile ring formation. **a.** Circumferential contractile stress acting around the division plane. The division plane is defined as normal to the division direction, 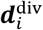, and passing through the centroid, 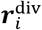. **b.** Circumferential contractile stress acting along the polarity direction in the division plane, ***p***_*j*(*i*)_. **c.** Polarity direction defined as the direction perpendicular to the division direction on the triangular plane, ***n***_*j*(*i*)_. **d.** Amplitude of contractile stress, 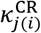 distributed in the division direction with a Gaussian centered on the division plane.

Contractile ring is formed by actin filaments aligned as a ring on the cell membrane, which generate active tensile stress to form a cleavage furrow (Fig. 3a). In this model, the stress oriented in the circumferential direction of each cell in the division plane is introduced into the surface of each triangle (Fig. 3b). By denoting the normal vector of the *j*th triangle of the *i*th cell as *n*_*j*(*i*)_, the orientation vector of the stress, represented by *p*_*j*(*i*)_, can be given by 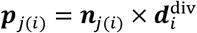 (Fig. 3c).

Anisotropic stress on each triangular surface is implemented by rewriting the area of the *j*th triangle in the *i*th cell as direction-dependent functions, represented by 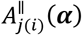 and 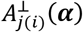, where ***α*** is a direction vector. In 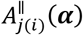 and 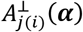, the perpendicular and parallel components of the position vector of each vertex to ***α*** are fixed as constants, respectively; that is, the changes in the area, represented by 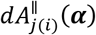 and 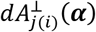, for the given displacement of a vertex corresponds to the changes in the area when the displacement has only parallel and perpendicular components to ***α***, respectively. The strict definition of 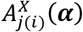 (*X*: an indicator of the component direction) is given in Appendix. Using this function, the energy of the anisotropic stress on the *j*th triangle surface in the *i*th cell, represented by *u*_*j*(*i*)_, can be described by separating it into two orthogonal components such that 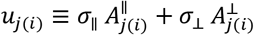, where the constants *σ*_∥_ and *σ*_⊥_ are the stresses exerted along the parallel and perpendicular directions, respectively [Eq. (A4) in Appendix]. Therefore, using *p*_*j*(*i*)_ and 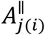 the energy function of the circumferential stress actively generated by the contractile ring of the *i*th cell, represented by 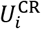, can be written by

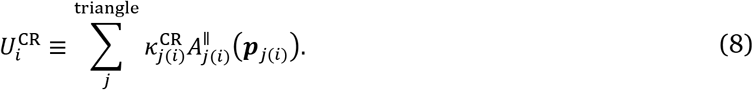

where constant 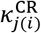 is the surface tension acting on the *j*th triangle of the *i*th cell.

Since the contractile ring is formed only during mitosis, the active stress of the contractile ring is set to zero during cell growth and a positive value after cell growth. Moreover, the magnitude of this stress is set to be distributed according to a Gaussian distribution along the division direction with the standard deviation of *l*_CR_ around the division plane (Fig. 3d). Therefore, by denoting the center position of the *j*th triangle of the *i*th cell as 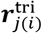, tension 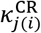 is given by

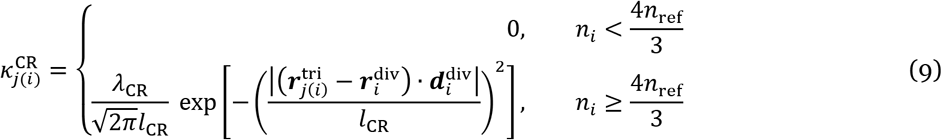

where constant *λ*_CR_ is the active tension of a contractile ring. Because constant 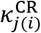 is given by time depending on the cell shape as in Eq. (9), the detailed balance is not satisfied. Therefore, the energy function of Eq. (8) generates active forces.

### 3.3 Description of division to daughter cells

The cell with the cleavage furrow is topologically divided into daughter cells (Fig. 2c-d). While active stress of the contractile ring in Eq. (8) gradually pinches off the cell, the triangular mesh expressing the cell membrane is rearranged by the topological operations described in the previous study [19]. In addition, when the open triangle is found in the cell (Fig. 4a), new two triangular surfaces are added at the open triangle (Fig. 4b). The positions of the added triangles are given as if the open triangles were completely overlapping. This topological operation splits one closed mesh into two separated meshes.

**Fig. 4:**
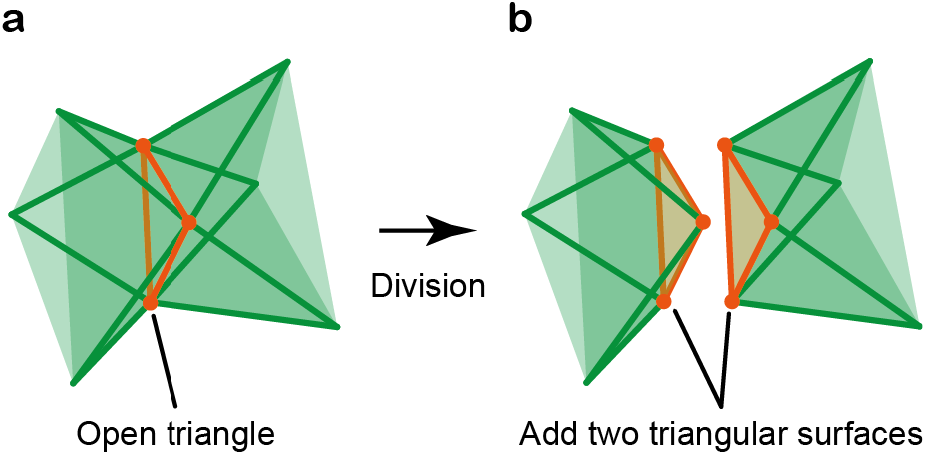
Topological operation of dividing a triangular mesh. A mesh expressing a single cell membrane is topologically divided at an open triangle (**a**) into two separated meshes expressing different cell membranes (**b**). The open triangle in each separated mesh is closed by adding a triangular surface. The locations of the added triangles are given as if the triangles completely overlap the open triangle.

## 4. Integrating the NCF and cell proliferation models for numerical simulations

To demonstrate the applicability of the proposed model, numerical simulations of proliferative cell dynamics were performed. For this purpose, in this section, the proposed model, described in Sec. 3, was integrated with the model of cell dynamics, described in Sec. 2. Cell behaviors, including cell proliferation, are described by energy and dissipation functions (Sec. 4.1), where the interactions between cells and substrate were described using the interfacial interaction model that we previously proposed [20]. This integrated model quantitatively describes mechanical behaviors of single-cell proliferation, such as the growth rate of cell volume and active stress of contractile ring (Sec. 4.2), and can be implemented for numerical simulations (Sec. 4.3).

### 4.1 Energy and dissipation functions of proliferative cell behaviors

To apply the proposed model to proliferating cell dynamics, *U* in Eq. (2) was expressed as

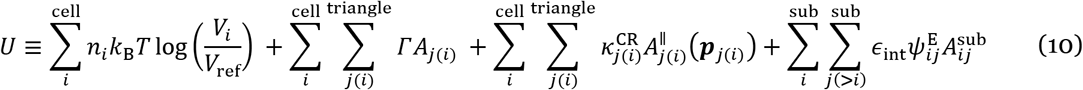

The first and second terms indicate the osmotic energy and cell surface energy described in Eq. (5). In the second term, the total surface area of the *i*th cell, 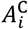, is described as the sum of the areas of all triangles comprising the cell, where the area of the *j* th triangle is represented by *A*_*j*(*i*)_. The third term indicates the energy of the contractile ring described in Eq. (8). The fourth term in Eq. (10) indicates the interfacial energy between cells and substrate, which is formulated in the previous study [20]. In this function, constant *ϵ*_int_ is the adhesion energy density, and 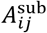 and 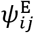 is the interfacial area and the interaction function of interfacial energy between the *i*th and *j*th subtriangular elements. Moreover, *W* in Eq. (4) was expressed as

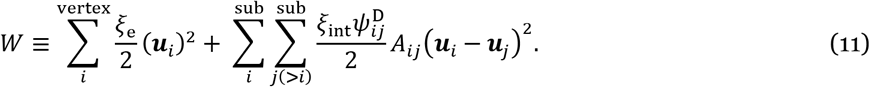

The first term in Eq. (11) indicates the viscous dissipation between the cell and the solvent, where constant *ξ*_e_ is the friction density coefficient. The second term indicates the viscous dissipation between cells and substrate, which is formulated in the previous study [20]. In this function, constant *ξ*_int_ is the friction density coefficient, and function 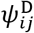 is the interaction function of interfacial dissipation between the *i*th and *j*th subtriangular elements. The details of the fourth term in Eq. (10) and the second term in Eq. (11), expressing interfacial interactions, are described in the previous study [20].

### 4.2 Non-dimensionalization and parameter setting

The regular triangle with the average area of each triangular mesh is considered, where the length of each edge composing the regular triangle is represented by *l*. Using *l*, the average vertex density per area is written by 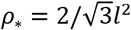, and the range of interface was set to *d*_0_ = *l*/2, the average length of the edges composing triangular elements for interfacial interaction. The physical parameters were non-dimensionalized by length *l*, time *τ* (= *μ/Γ*), and energy *Γl*^2^, by fixing the values of *l, μ*, and *Γ*. Physical parameters used for the simulations are listed in Table 1.

**Table 1:** Physical parameters for simulations.

| Parameter | Unit | Explanation |
| --- | --- | --- |
| $l$ | - | Edge length (set to be a unit) |
| $\Gamma$ | - | Cortical tension of cell membrane (set to be a unit) |
| $\mu$ | - | Planar viscosity of cell membrane (set to be a unit) |
| $L_x, L_y, L_z$ | $l$ | System box size (= 50, 52, 100) |
| $V_{\text{ref}}$ | $l^3$ | Reference cell volume (= $10^3$) |
| $\xi_e$ | $\mu$ | Friction density from solvent (= $10^{-3}$) |
| $\lambda_{\text{CR}}$ | $\Gamma l$ | Tension of contractile ring (= 30) |
| $l_{\text{CR}}$ | $l$ | Standard deviation of contractile ring (= 0.6) |
| $\tau_{\text{cc}}$ | $\tau$ (= $\mu/\Gamma$) | Cell cycle period (= 60) |
| $\epsilon_{\text{int}}$ | $\Gamma$ | Interfacial energy density (= 1) |
| $\xi_{\text{int}}$ | $\mu$ | Interfacial friction density (= 1) |

The typical size of Hela and MDA MB231 cells is about 15~17μm [22], from which the cell volume is estimated to be about *V*_ref_ = 4×10^3^μm^3^. By fixing *V*_ref_ = 10^3^*l*^3^, the unit length is set to *l* = 1.6μm.

Cortical tension widely varies with cell types and states in 30~4000pN/μm (summarized in [23]); for example, HeLa cells increase their surface tension from 200 pN/μm during interphase to 1,600 pN/μm during metaphase [24], and an active tension of S180 cells is estimated as 400 pN/μm[25]. Therefore, the surface energy density, corresponding to the unit energy density, was set to *Γ* = 400 pN/μm.

Elastic modulus and the effective thickness of actin cortex, represented by *E*_crt_ and *t*_crt_, are estimated as 15 kPa and 0.3 μm, respectively, regarding S180 cells [25]. Moreover, the lifetime of an actin cortex, represented by *τ*_crt_, is typically a few tens of seconds [26,27]. Let us consider an actin cortex that extends with a mean strain velocity, 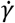, then the mean strain of the cortex can be written by 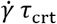. Hence, the stress exerted on the cortex, represented by *σ*, can be written by 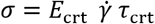, from which the effective viscous coefficient of the cortex can be estimated as *E*_crt_ *τ*_crt_. By considering the thickness of the cortex, we obtained the planar viscosity as *μ* = *E*_crt_ *τ*_crt_ *t*_crt_ (= 135 × 10^3^ pN s/μm). Therefore, the unit time is given as *τ* (= *μ/Γ*) ≈ 5 min.

The main component of the solvent for cells is water, whose viscosity is 0.69 pNs/μm^2^, from which the friction density coefficient of a triangular element with the length size of *l* in the solvent was approximately set to *ξ*_e_ = 1 pNs/μm (10^-3^ μ), much lower than the other viscous effects. Moreover, the adhesion energy density can be estimated as *ϵ*_int_ = 0.16~1200 pN/μm (0.0016~12 *Γ*) [20]. Hence, by considering the balance of the adhesion with the cell surface tension, the adhesion energy density was set to *ϵ*_int_ = *Γ*. Moreover, the timescale of cell adhesion is of the order of a few seconds, which gives an expected friction density of *ξ*_int_ = *μ*.

The width of the cortical ring varies with cell types and ranges from 0.13 to 8 μm [1]. Considering the 95% range of the distribution function in Eq. (9) as the width of the contractile ring, for example, HeLa cells have a width of 4 μm, yielding *l*_CR_ = 0.6 *l*. Moreover, the constriction duration of the contractile ring, represented by *τ*_CR_, also varies with cell types from 5 to 34 min [1], yielding *τ*_CR_ = 11 min. Given that the initial perimeter length of the contractile ring expressed as *P*_CR_ is approximately 30π μm, the mean rate of the constriction can be roughly estimated as *P*_CR_/*τ*_CR_. Using the planar viscosity of actin cortex, *μ*, the active tension of the contractile ring can be written as *λ*_CR_ = *μ* (*P*_CR_/*τ*_CR_) and estimated as 30 *Γl* (= 1.9 × 10^4^ pN). Furthermore, the cell cycle ranges widely from 30 min to days, and as an example, it was set to *τ*_cc_ = 60 *τ* (=5 hr).

### 4.3 Numerical calculations

To perform numerical simulations, the time development of vertex locations and remeshing are alternately calculated. Vertex velocities, *u_i_*, are calculated by solving the simultaneous equations of Eq. (4) for all vertices. The time development of the vertex locations, ***r**_i_*, is calculated by numerically integrating Eq. (4) using the Runge–Kutta method with a time step Δ*t* (= 2.0 × 10^-3^*τ*). The details of numerical procedures are similar to those described in the previous study [20], and the numerical parameters used for the simulations are listed in Table 2. The numerical algorithm was implemented in C++ programming language, where OpenMP was used for parallel computations. All experiments were performed on workstations with 3.2 GHz Intel Xeon dual processors and 64 GB RAM.

**Table 2:** Numerical parameters for simulations.

| Parameter | Unit | Explanation |
| --- | --- | --- |
| $\Delta t$ | $\tau$ | Time step of vertex displacements (= $2.0 \times 10^{-3}$) |
| $\Delta t_{\text{R}}$ | $\tau$ | Time step of topological operations (= $10^{-2}$) |
| $d_0^\perp$ | $l$ | Depth of interface (= 1/2) |
| $d_0^\parallel$ | $l$ | Width of interface (= 1/2) |
| $\Delta_{\text{max}}$ | $l$ | Maximum value of drift (= $10^{-2}$) |
| $\Delta_{\text{th}}$ | $l$ | Drift threshold (= $10^{-3}$) |

## 5. Numerical simulations of proliferative multicellular dynamics

The novelty of the proposed model lies in the detailed flexible description of the process of cell proliferation in multicellular systems. To demonstrate the applicability of the proposed model, in this section, we examined cell dynamics in three distinct situations. First, to clarify that the model can recapitulate the process of cell proliferation, we observed the detailed process of a single cell proliferation (Sec. 3.1). Second, to show that the model can be applied to analyze the process of cell proliferation, we investigated effects of the actomyosin orientation of contractile ring on the cleavage formation (Sec. 3.2). Third, to show the applicability of the proposed model to tissue-scale dynamics, we investigated multicellular dynamics emerging from successive cell proliferation on a substrate (Sec. 3.2).

### 5.1 Recapitulation of the cell proliferation process

To demonstrate that the proposed model can describe the detailed process of cell proliferation, numerical simulations of single-cell proliferation were performed. In the *xyz*-orthogonal coordinates, a rectangular system box was considered within 0 ≤ *x* < *L_x_*, 0 ≤ *y* < *L_y_*, and 0 ≤ *z* < *L_z_*. As an initial condition, a spherical cell with a number of molecules of *n*_ref_ is assumed to be in a stationary solvent. The cell was expressed by a triangular mesh, which was pre-randomized and optimized before the simulations. The orientation of the active stress in the contractile ring was set to be circumferential in the division plane (Fig. 4a). Division direction, 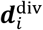, was set to be along the *z*-axis. Simulations were performed during the time range of 80τ.

Numerical simulations showed that our model can simulate the detailed process of single-cell proliferation (Fig. 5, Supplementary Video 1). The cell size gradually increased during the growth phase (0 ≤ *t* ≤ 30*τ*), after which a contractile ring was formed (*t* = 30*τ*). The contractile ring shrunk the center of the cell and formed a cleavage furrow (30*τ* ≤ *t* ≤ 33*τ*), at which the cell was divided to daughter cells (*t* = 33*τ*). Following the division, the daughter cells adhered to each other (34*τ* ≤ *t* ≤ 36*τ*). The cell volume increased linearly with time during the growth phase (0 ≤ *t* ≤ 30*τ*), and abruptly decreased by the contractile ring formation (*t* = 30*τ*) (Fig. 6a). To further evaluate the temporal changes in cell shape, the sphericity of a cell was defined as the ratio of the radii of spheres with the same volume and surface area. The cell sphericity remained high during the growth phase, transiently decreased after the contractile ring formation, and recovered after the division (Fig. 6b). These results show that the proposed model successfully described the detailed process of cell proliferation, i.e., volume growth, contractile ring formation, and division to daughter cells.

**Fig. 5:**
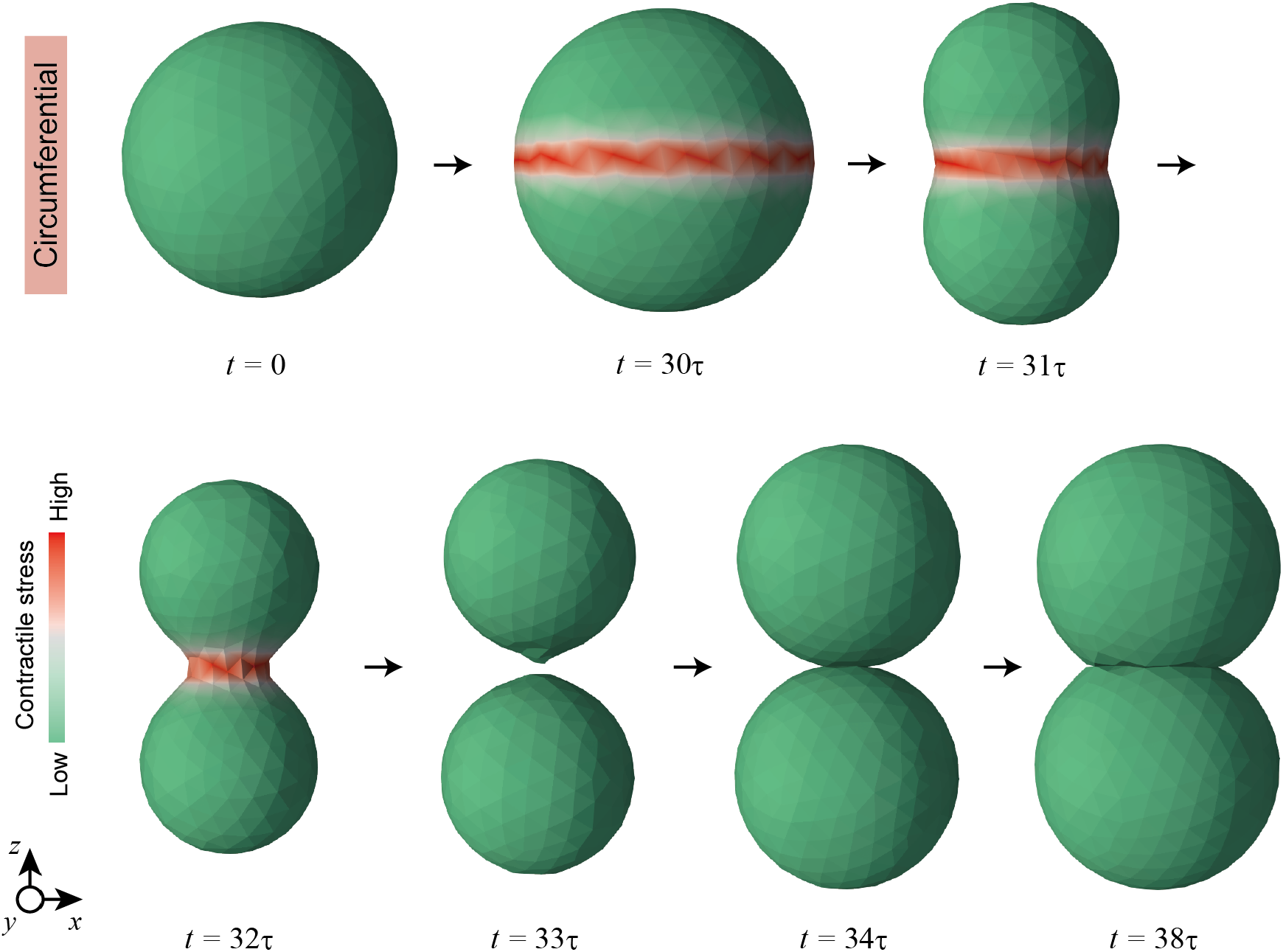
Numerical simulation of single-cell proliferation process. Time series of cell shapes resulting in the case with circumferential contractile stress. Amplitude of contractile stress, 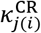, is colored with a slope from green to red. This process is also shown in Supplementary Video 1.

**Fig. 6:**
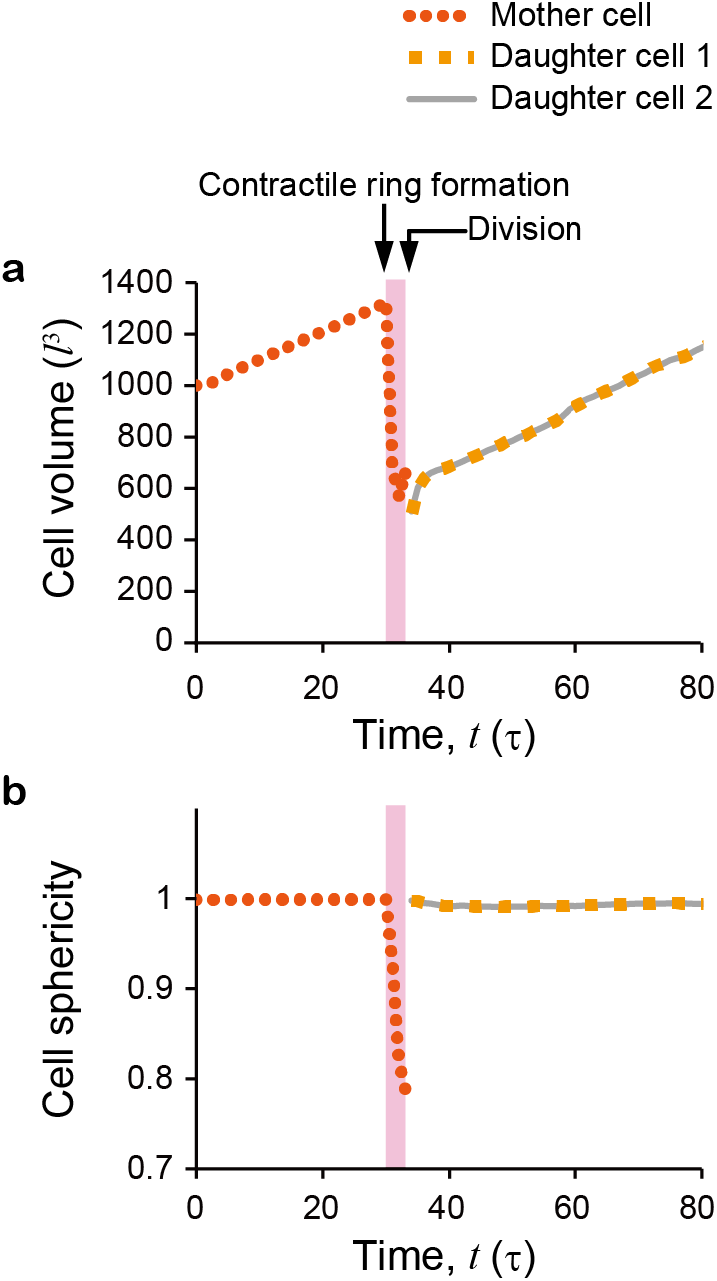
Quantifications of proliferating cell morphologies. **a, b.** Cell volume and sphericity as functions of time during the proliferation process, respectively. One mother cell (orange dotted line) divides into two daughter cells (yellow dashed line and gray solid line). The period colored pink indicates the period during which the contractile ring is formed. These results were obtained from the simulation shown in Fig. 5.

### 5.2 Role of actomyosin stress orientation in cleavage furrow formation

To demonstrate that the proposed model can account for various regulations of cell proliferation, as an example, we investigated the effects of the orientation of active stress in the contractile ring on cleavage formation. Here, we considered two situations, i.e., circumferential stress (Fig. 3b) and isotropic stress (Fig. 7a). While the circumferential stress is described through the energy function of Eq. (8), the isotropic stress is introduced via an energy function, given by

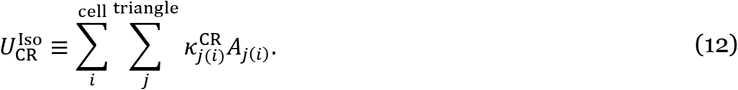

**Fig. 7:**
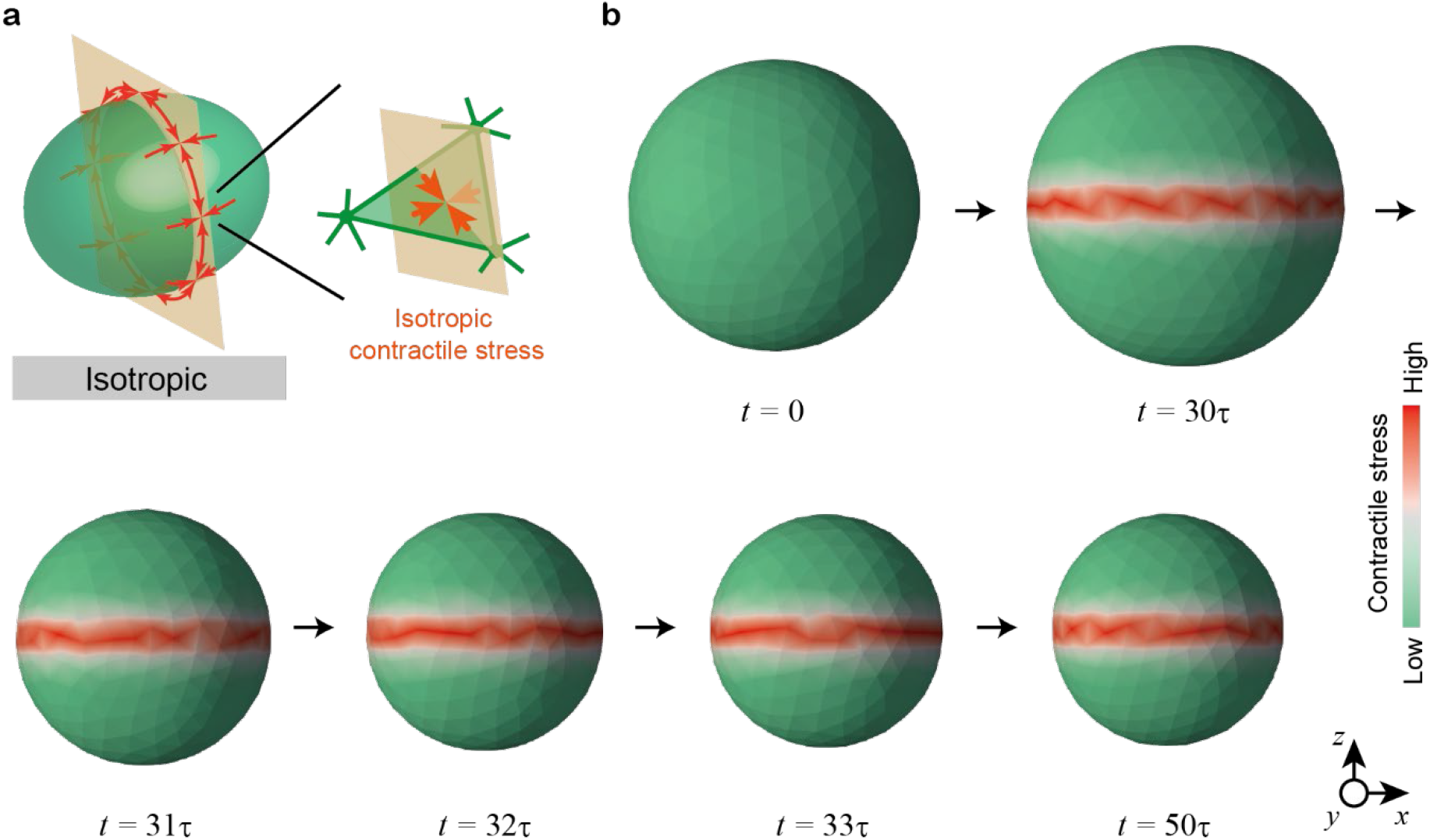
Effect of isotropic stress of contractile ring. **a.** Schematic illustration of isotropic contractile stress acting around the division plane. **b.** Time series of cell shapes resulting in the case with isotropic contractile stress. In **b**, amplitude of contractile stress, 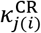, is colored with a slope from green to red. This process is also shown in Supplementary Video 2.

In the situation with isotropic stress, the third term on the right-hand side of Eq. (10) replaced Eq. (12). In both situations, the division direction, 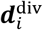, is set to be along the *z*-axis.

As a result, while the circumferential stress caused the cell to locally contract at the division plane to form a cleavage furrow (Fig. 5), the isotropic stress could not (Fig. 7b, Supplementary Video 2). The isotropic stress caused the cell to entirely contract while maintaining the cell shape spherical (30*τ* ≤ *t* ≤ 32*τ*). No cleavage furrow was formed, thus maintaining a contractile ring and the resulting cell contraction (32*τ* ≤ *t* ≤ 50*τ*).

The ability to form a cleavage furrow lies on the summation forces acting on cell membranes, which differ between the cases with circumferential and isotropic stresses (Fig. 8). Circumferential stress mainly generates forces in the radial direction, squeezing the cell in the dividing plane (Fig. 8a). On the other hand, isotropic stress mainly provides the force contracting the cell in the direction of the division axis (Fig. 8b). Quantitatively, both the radial and axial forces are produced in both the cases of circumferential and isotropic stresses, albeit with a quite difference in their balance (Fig. 8c, d). In the case with circumferential stress (red curves), the radial force is ca. five times larger than the axial force. In the case with isotropic stress, the radial force is ca. four times smaller than the axial force (gray curves). The small axial force in the case with circumferential stress (Fig. 8d) is due to the membrane surrounding the furrow drawn into the contractile ring.

**Fig. 8:**
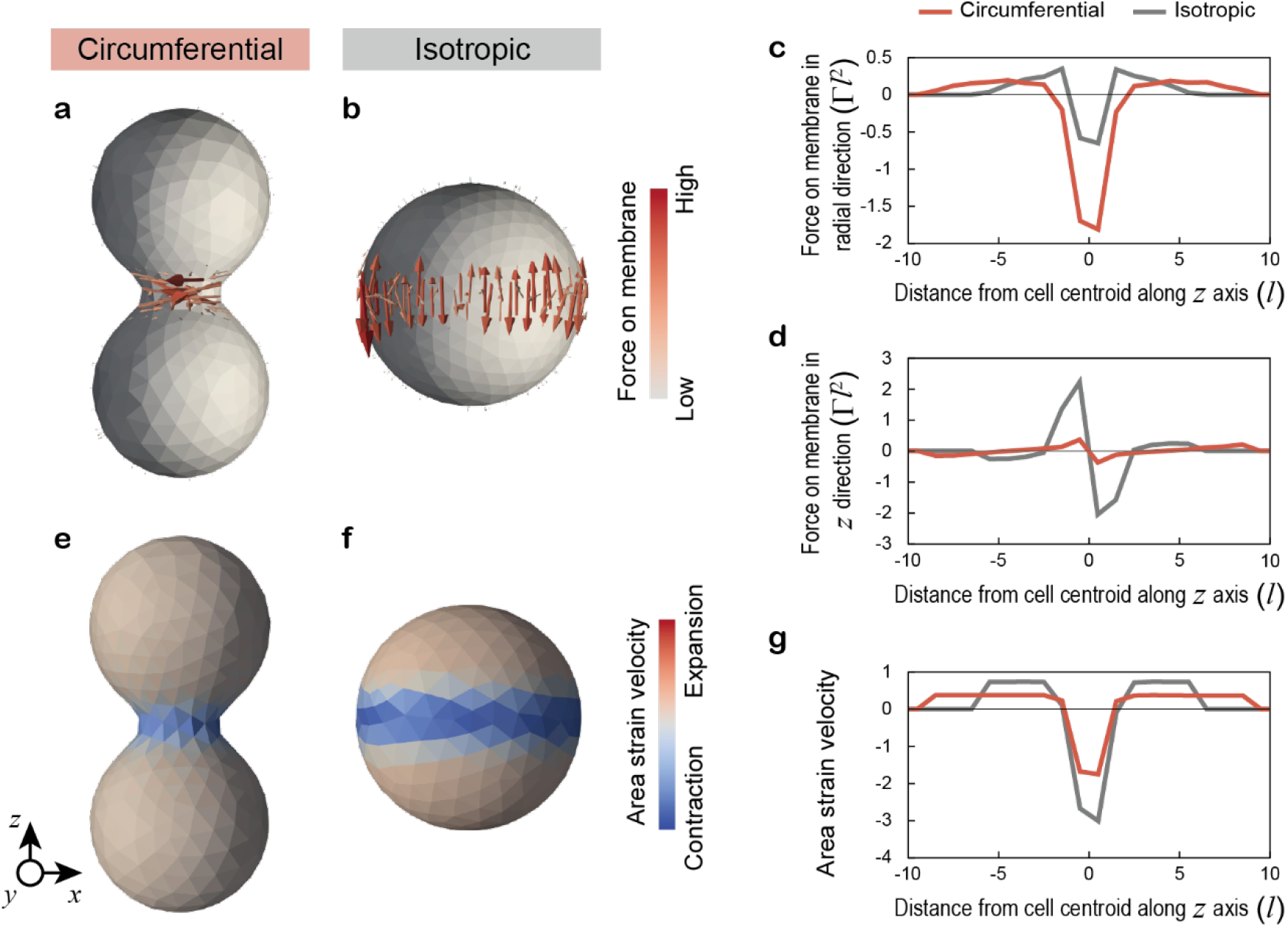
Cell membrane dynamics induced by contractile ring. **a, b.** Distributions of summation forces exerted on the cell membrane in cases with circumferential and isotropic contractile stresses, respectively. Cells in **a, b** are truncated in the *xz* plane through their centroids. **c, d**. Forces on the cell membrane in the radial and *z* directions, respectively. **e, f.** Distributions of area strain velocity of the cell membrane, 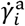, in cases with circumferential and isotropic contractile stresses, respectively. **g**. Spatial profiles of area strain velocity in the *z* direction. In **c, d**, and **g**, Each profile is calculated as the average as a function of the distance from the cell centroid. The results were obtained from the simulations at *t* = 32*τ* shown in Figs. 5 and 7.

In our model, the formation of a cleavage furrow requires circumferential stress, while in a previous study [18], isotropic stress was found to be sufficient for cleavage furrow formation. This discrepancy may be attributed to the differences in the properties of cell surface membranes considered in the two models. Specifically, while the previous study regarded the membrane as an elastic body, the NCF model takes into account the non-conservative nature of the membrane [19]. To evaluate the contribution of the nonconservative feature to the cleavage formation, the area strain velocity of the membrane at the *i*th triangle, represented by 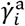, was calculated by

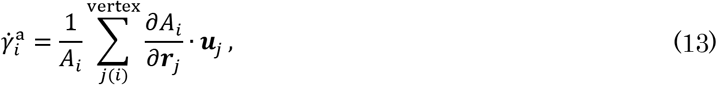

where the summation is for all vertices composing the *i*th triangle. As a result, in case of both the circumferential and isotropic stresses, the cell membranes contract around the division plane (Fig. 8e, f). However, their strain is slight in the case of circumferential stress and large in the case of isotropic stress (Fig. 8g). Specifically, circumferential stress globally contracts the perimeter of a cell, leading to the formation of a cleavage furrow. In contrast, isotropic stress locally reduces the membrane area around the division plane, flattening the membrane and suppressing the formation of a cleavage furrow. These results suggest that the circumferential orientation of active stress in the contractile ring plays a crucial role in cleavage furrow formation.

### 5.3 Replication of cell sheet formation and stratification on substrate

To demonstrate the applicability of the model to proliferative multicellular dynamics, numerical simulations of tissue growth on a substrate were performed. The orientation of actomyosin stress of the contractile ring was set to be circumferential in the division plane as described in Eq. (8), and the total energy function was given by Eq. (10). Division direction, 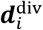, was set to be normal to the substrate plane (i.e., along the *z*-axis) as well as along the longest axis of each cell.

A rectangular system box was considered within 0 ≤ *x* < *L_x_*, 0 ≤ *y* < *L_y_*, and 0 ≤ *z* < *L_z_* in the orthogonal *xyz*-coordinates. Periodic boundary conditions were applied to all boundaries. As an initial condition, a spherical cell with a number of molecules of *n*_ref_ is assumed to be in a stationary solvent. A solid substrate is inserted in the *xy*-plane with a slight gap to the bottom surface of the cell. Both the solid substrate and the cell were expressed by a triangular mesh, which was pre-randomized and optimized before the simulations. During simulations, the vertex positions and mesh topology of the substrate were fixed on the coordinates. Simulations were performed during the time range of 450*τ*.

Numerical simulations showed that the model successfully recapitulated the process of tissue growth driven by successive cell proliferations at the single-cell resolution (Fig. 9a, Supplementary Video 3). In this process, a single cell grew and divided into multiple cells, and spread over the substrate (0 ≤ *t* ≲ 330*τ*). After almost covering the substrate, the cells further grew and divided to form a stratified structure (330*τ* ≲ *t* ≲ 450*τ*). During the tissue growth, individual cells tightly adhere to each other to form continuous boundary surfaces (Fig. 9b, Supplementary Video 4). When cells form a monolayer, each cell has a polygonal shape in the planar direction of the substrate and has a plump, rounded shape on the tissue surface (Fig. 9c). Therefore, this model enables us to analyze multicellular dynamics in the growing tissue composed of more than a hundred cells with successive cell proliferation. Notably, the proposed model also allowed a detailed observation of the proliferation process of each cell embedded in the tissue with more than a hundred cells (Fig. 9d, Supplementary Video 5). For example, the increase in the volume of a particular cell (300*τ* ≲ *t* ≲ 354*τ*), contractile ring formation (355*τ* ≲ *t* ≲ 360*τ*), division to daughter cells (*t* ≈ 362*τ*), and adhesion between the cells (362*τ* ≲ *t* ≲ 400*τ*) could be successfully observed. The morphologies observed in this cell were significantly different from those of the single cell (Fig. 5), due to the effects of adhesion to the surrounding cells and substrate.

**Fig. 9:**
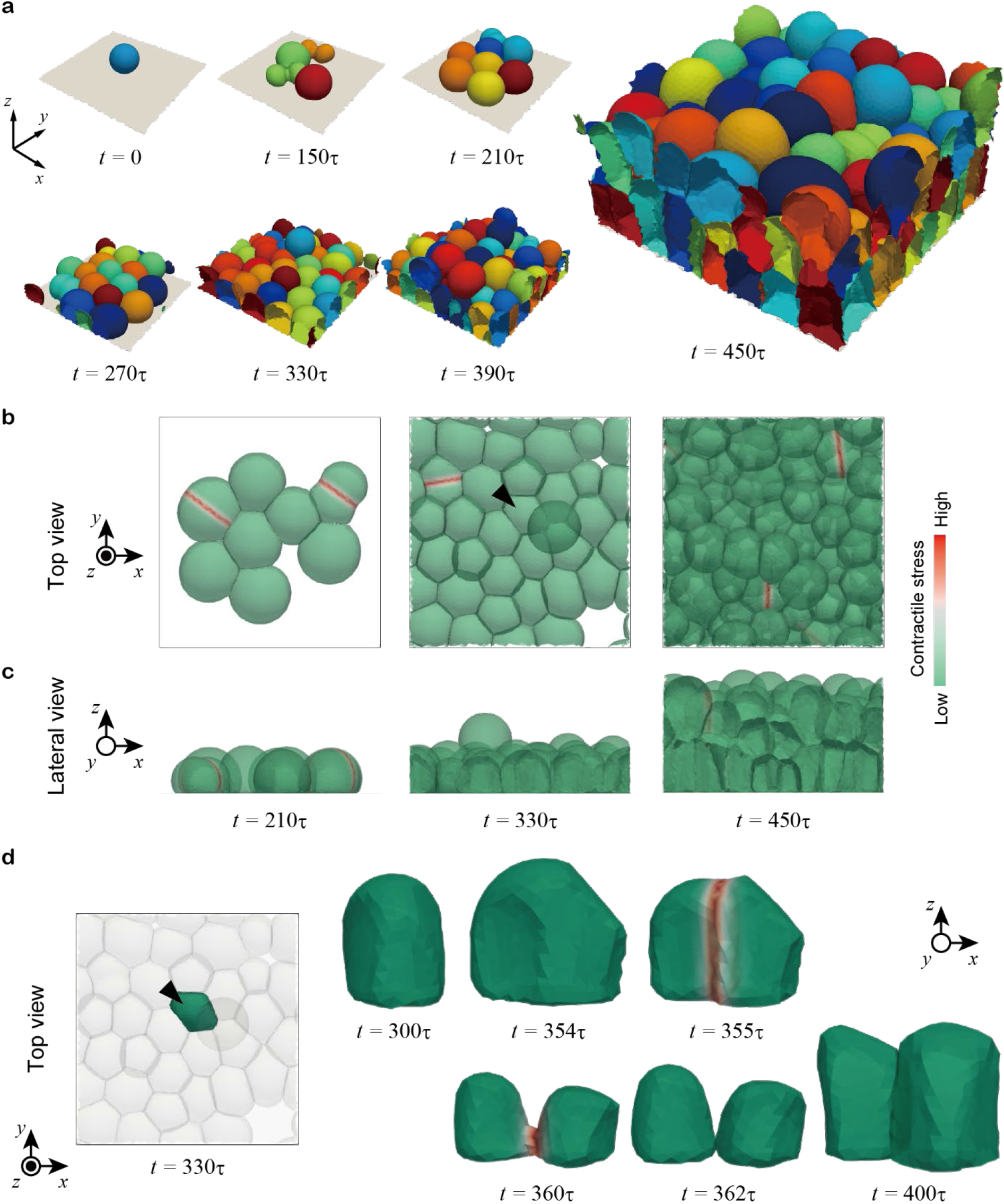
Numerical simulations of tissue growth on substrate. **a.** Time series of proliferating multicellular dynamics. Individual cells are shown in different colors. **b, c.** Top and lateral views of the time series of proliferating multicellular dynamics, respectively. **d.** Time series of the proliferation of a single cell embedded in the tissue. The observed cell is those pointed by arrows in **b** and **d**. In **b**-**d**, amplitude of contractile stress, 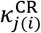, is colored with a slope from green to red. The processes in **a, b**, and **d** are also shown in Supplementary Videos 3, 4, and 5, respectively.

Figure 10 shows the physical quantities of the growing tissue. The total number of cells increased exponentially over time (Fig. 10a). However, the growth rate is slower than the theoretical solution due to the model not accounting for the time period of cell contraction caused by the contractile ring. The average cell volume and sphericity exhibited periodic fluctuations in response to the cell cycle (Fig. 10b, c). Because the cell growth rate was set to vary, the timing of contractile ring formation shifted as the tissue grew, and the amplitudes of changes in cell volume and cell sphericity decreased over time. Additionally, the average values of cell volume and cell sphericity decreased gradually as cells were densely packed and compressed. It is noteworthy that these behaviors were not observed in the single-cell proliferation process (Fig. 5) but resulted from the adhesive interactions between cells and substrate. To account for these interactions, the model of cell adhesion [20], which is an integral component of the proposed model plays a key role. Overall, the proposed model enables quantitative analyses of proliferative multicellular dynamics on multiple scales, including the detailed processes of single-cell proliferation, resulting interactions with surrounding cells and substrate, and the entire tissue dynamics.

**Fig. 10:**
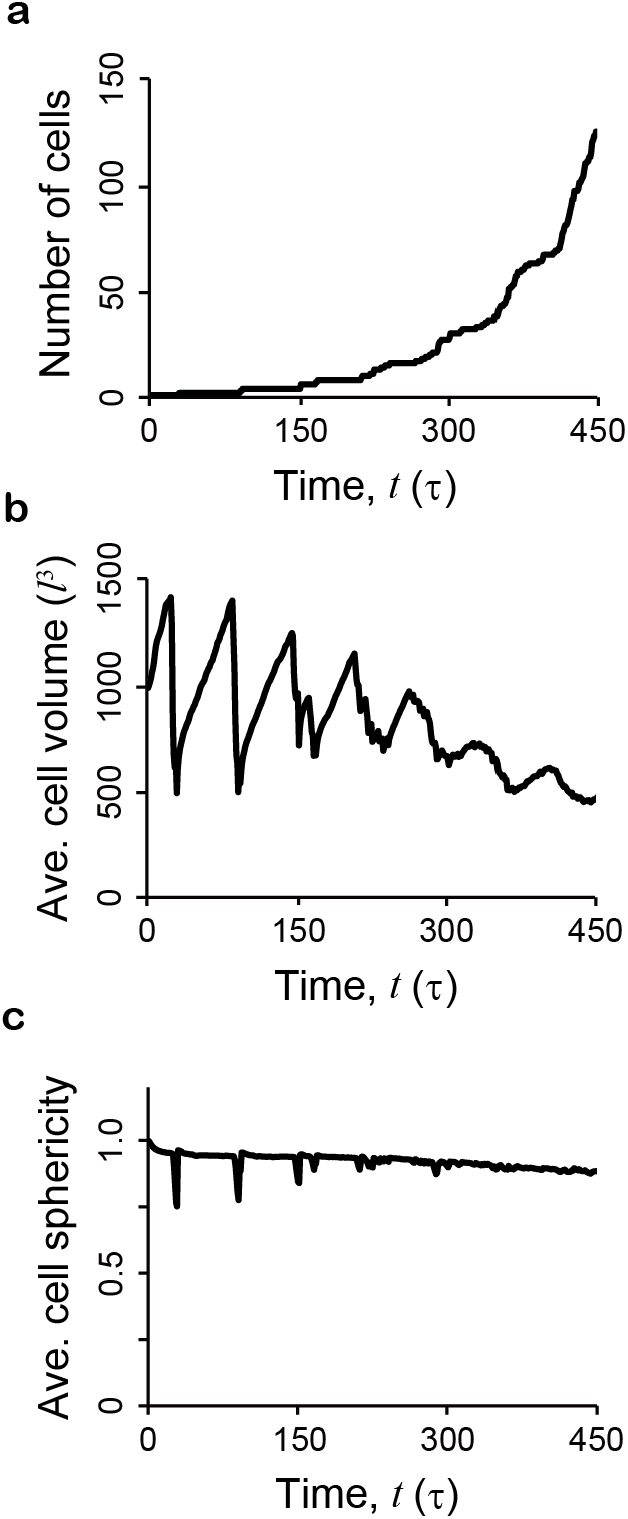
Quantifications of proliferating cell morphologies. **a-c.** Number of cells, average cell volume, and average cell sphericity as functions of time, respectively. These results were obtained from the simulation shown in Fig. 9.

## 6. Discussion

A new model of cell proliferation based on the NCF model was proposed for simulating long-term multicellular dynamics accompanied by successive cell proliferation. The proposed model recapitulated individual-cell proliferations: i.e., cell volume growth, cleavage furrow formation, and division to daughter cells in the subcellular resolution. For example, numerical simulations suggested that active stress of the contractile ring can form a cleavage furrow if it is circumferential, but not if it is isotropic, indicating that the orientation of actomyosin in the contractile ring is important for cells to form cleavage furrows. A further simulation successfully recapitulated successive proliferation of adhesive cells on a substrate, causing cell a sheet formation and stratification, where the process of each cell proliferation occurring in a hundred of proliferating cells can be observed. These results showed that the proposed model enables us to analyze tissue-scale multicellular dynamics at the subcellular resolution.

This model has several advantages: it can express the process of volume growth, cleavage furrow formation, and division to daughter cells for each cell. These behaviors are described by effective energy and dissipative functions, which can be flexibly modified for specific biological situations. This simple description enables us to analyze the effects of various cell proliferation behaviors on multicellular dynamics, e.g., cell growth rate, direction and location of the contractile ring, and orientation and amplitude of actomyosin contraction. Moreover, while this model can express the proliferation process of each cell, it can be applied to the dynamics of the entire tissue composed of more than a hundred cells. That is, the proposed model enables us to analyze tissue-scale mechanical interactions among proliferative cells in the subcellular resolution. This comprehensive description distinguishes the proposed model from other models of proliferative multicellular dynamics that totally ignore the process of cleavage furrow formation by a contractile ring and simply describe the mitotic process by dividing each cellular object, such as vertex models [14,28] and cell membrane models [15,16].

The proposed model presents opportunities for further refinement and development. Specifically, as cytoskeletal dynamics are crucial in cleavage formation processes [29], explicitly incorporating cytoskeletal dynamics independently of cell membrane dynamics would improve this model. For instance, cortical actins, which line the lipid membrane, are gradually localized during the formation of the contractile ring, a process not accounted for in the current model. This localization can be expressed by defining quantities of local states on the membrane and remapping their distributions at each remeshing occurrence [28]. Moreover, spindle dynamics play a vital role in determining the location and orientation of the division plane. The effects of structural components beyond cortical actins, such as microtubules, could also be incorporated through the external force term in this model. These enhancements will enable the simulation of proliferative multicellular dynamics in realistic scenarios, including mitotic rounding and pinching processes, asymmetric divisions, and their interactions with surrounding cells and substrates.

## 7. Conclusion

Cell proliferation plays key roles in embryogenesis, homeostasis, wound healing, and carcinogenesis. The process of cell proliferation involves cell growth, contractile ring formation, and division to daughter cells, causing geometrical and mechanical effects on the surrounding cell population. To address proliferative multicellular dynamics, this paper reports on the model of cell proliferation based on the NCF model framework. The proposed model successfully recapitulated the proliferation process of each cell in the subcellular resolution, enabling us to analyze the effects of various cell proliferation behaviors on multicellular dynamics, e.g., cell growth rate, direction and location of contractile ring, and orientation and amplitude of actomyosin contraction. The proposed model can be applied to tissue-scale multicellular dynamics such as epithelial tissue formation and stratifications from cytoskeletal dynamics and cell-cell and cell-substrate interactions. Thus, the proposed framework provides a useful basis for analyzing a wide range of proliferative cell dynamics and will lead to a new approach for understanding the mechanics of 3D multicellular systems.

## Appendix

### A1. Energy function for inducing anisotropic stress on triangular surface

Anisotropic stress exerted on an arbitrary triangle in the *xyz*-orthogonal coordinates is considered. The triangle is composed of vertices 0, 1, and 2. Normal vector of the triangle is represented by ***n*** in the *xyz*-orthogonal coordinates. In addition, the *x′y′* orthogonal coordinates can be locally defined in the plane of the triangle, whose basis vectors are represented by ***e**_x′_* and ***e**_y′_* (= ***n × e***_*x′*_). Using ***e**_x′_* and ***e**_y′_*, the area of the triangle, represented by *A*, is given by

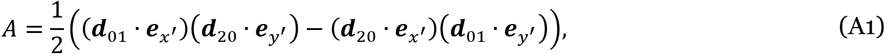

where ***d**_αβ_* is the distance vector from the *α*- to *β*- vertices.

By regarding the components of vertex locations along ***e**_y′_* as constants in Eq. (A1), Eq. (A1) is rewritten as follows

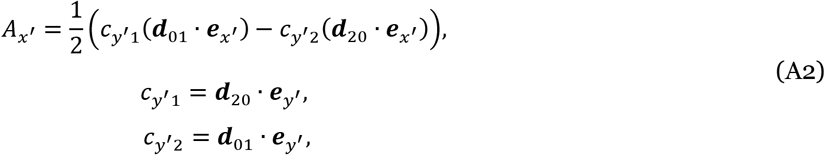

where *c*_*y′*1_ and *c*_*y′*2_ are regarded as constants. Similarly, by regarding the components of vertex locations along ***e**_x′_* as constants in (A2), Eq. (A1) is given by

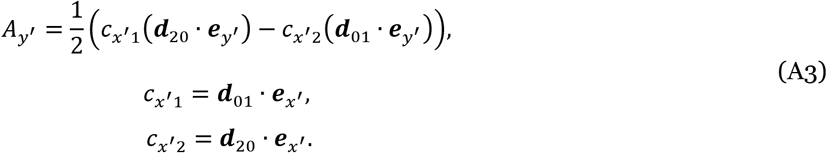

where *c*_*x′*1_ and *c*_*x′*2_ are regarded as constants.

Using Eqs. (A2) and (A3), the energy of anisotropic stress in the triangle surface, represented by *u*, can be given by

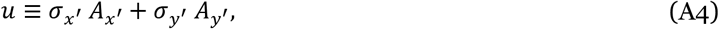

where the constants *σ_x′_* and *σ_y′_* are the exerted stress along ***e**_x′_* and ***e**_y′_*, respectively. When the exerted tension is isotropic (*σ_x′_* = *σ_y′_* = *σ*/2), the energy function becomes *u* = *σA*.

By calculating the partial derivatives of Eq. (A4), the forces acting on vertices 0, 1, and 2, represented by ***f***_0_, ***f***_1_, and ***f***_2_, respectively, were given by

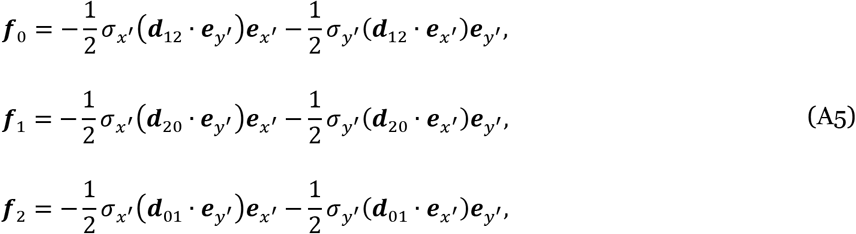

respectively. The sum of each of the forces and moments in Eq. (A5) becomes zero. According to the above formulation, the area of the *j*th triangle in the *i*th cell is rewritten as a direction-dependent function, where the components of vertex position vectors in the direction normal to *α* is regarded as constants, 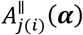.

## Supplementary Materials

**Video 1. Process of cell growth, contractile ring formation, and division to daughter cells**

The amplitude of contractile stress is colored with a slope from green to red. The snapshots of this process are shown in Fig. 5.

**Video 2. Effects of isotropic stress of contractile ring on cleavage furrow formation**

The amplitude of contractile stress is colored with a slope from green to red. The snapshots of this process are shown in Fig. 7.

**Video 3. Process of tissue growth by successive cell proliferation on substrate**

Individual cells are shown in different colors. The snapshots of this process are shown in Fig. 9a.

**Video 4. Top-view of tissue growth process by successive cell proliferations on substrate**

The amplitude of contractile stress is colored with a slope from green to red. The snapshots of this process are shown in Fig. 9b.

**Video 5. Proliferation process of a single cell embedded in growing tissue**

The amplitude of contractile stress is colored with a slope from green to red. The snapshots of this process are shown in Fig. 9d.

## Author Contribution

SO conceived the project, developed the software, and performed the analyses; TH provided ideas; SO wrote the first draft of the manuscript, and SO and TH reviewed and edited. All authors contributed to the final manuscript.

## Acknowledgment

This work was supported by the Japan Science and Technology Agency (JST), CREST Grant No. JPMJCR1921; the Japan Agency for Medical Research and Development (AMED), Grant No. 21bm0704065h0002; the Japan Society for the Promotion of Science (JSPS), KAKENHI Grants No. 21H01209, 22K18749, and 22H05170; the World Premier International Research Center Initiative, Ministry of Education, Culture, Sports, Science and Technology (MEXT), Japan (to SO); and the seed grant of Mechanobiology Institute (to TH).

---

**Supplementary Fig. 1.**
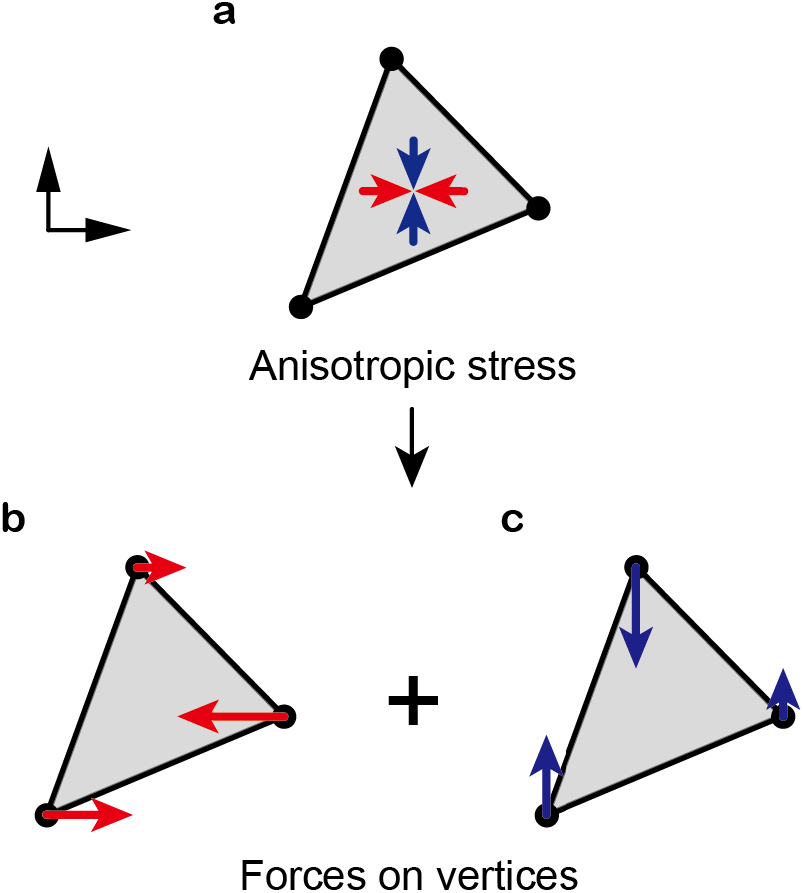
Description of anisotropic stress on a triangular element. Anisotropic stress acting on a triangle is expressed as a set of forces applied to vertices in two mutually orthogonal directions.

